# A glucose-dependent spatial patterning of exocytosis in human β-cells is disrupted in type 2 diabetes

**DOI:** 10.1101/534669

**Authors:** Jianyang Fu, John Maringa Githaka, Xiaoqing Dai, Gregory Plummer, Kunimasa Suzuki, Aliya F. Spigelman, Austin Bautista, Ryekjang Kim, Dafna Greitzer-Antes, Jocelyn E. Manning Fox, Herbert Y. Gaisano, Patrick E. MacDonald

**Affiliations:** Alberta Diabetes Institute and the Department of Pharmacology, University of Alberta, Edmonton, AB, T6G 2E1, Canada; Department of Biochemistry, University of Alberta, Edmonton, AB, T6G 2E1, Canada; Departments of Medicine and Physiology, University of Toronto, Toronto, ON, M5S 1A8, Canada

**Keywords:** diabetes, islets of Langerhans, insulin, exocytosis, ion channels, SUMOylation

## Abstract

Impaired insulin secretion in type 2 diabetes (T2D) is linked to reduced insulin granule docking, disorganization of the exocytotic site, and an impaired glucose-dependent facilitation of insulin exocytosis. We show in β-cells from 80 human donors that the glucose-dependent amplification of exocytosis is disrupted in T2D. Spatial analyses of granule fusion, visualized by total internal reflection fluorescence (TIRF) microscopy in 24 of these donors, demonstrate that these are non-random across the surface of β-cells from donors with no diabetes (ND). The compartmentalization of events occurs within regions defined by concurrent or recent membrane-resident secretory granules. This organization, and the number of membrane-associated granules, is glucose-dependent and notably impaired in T2D β-cells. Mechanistically, multi-channel Kv2.1 clusters contribute to maintaining the density of membrane-resident granules and the number of fusion ‘hotspots’, while SUMOylation sites at the channel N-(K145) and C-terminus (K470) determine the relative proportion of fusion events occurring within these regions. Thus, a glucose-dependent compartmentalization of fusion, regulated in part by a structural role for Kv2.1, is disrupted in β-cells from donors with type 2 diabetes.

**HIGHLIGHTS:**

- Exocytosis of secretory granules is non-random across the surface of human b-cells, and this organization is disrupted in type 2 diabetes.
- Increasing glucose facilitates the spatial compartmentalization of fusion, independent of an overall increase in event frequency.
- Compartmentalized ‘hotspots’ occur at sites marked by membrane-associated granules, the density of which is regulated in part by a clustered K+ channel (Kv2.1).
- SUMOylation status of the channel controls the proportion of events that occur within these local regions.

## INTRODUCTION

Insulin, secreted from pancreatic islet β-cells, is a critical regulator of blood glucose and energy homeostasis in humans (1). Impaired insulin secretion is a hallmark of type 2 diabetes (T2D) which results from a complex interplay between reduced insulin sensitivity and impaired insulin secretion (2). A tipping point occurs when secretion from genetically predisposed β-cells fails to meet the demands of peripheral insulin resistance (3, 4). While impaired insulin responses in T2D may result from a long-term reduction in β-cell mass (5), reduced secretory function likely predominates at earlier stages (6, 7). In isolated islets or β-cells from donors with T2D in vitro, exocytotic protein expression is reduced (8, 9) and exocytotic responses are impaired (10, 11). Most recently this has been attributed to an impaired glucose-dependent granule docking and reduced expression of docking and related active zone proteins (12).

While β-cells lack ultra-structurally identifiable active zones, glucose-stimulation *in situ* triggers fusion events that are progressively localized to preferential release sites (13). These appear directed towards the islet vasculature (13–15) and are regulated by extracellular matrix (ECM) interactions with integrin receptors (16), which likely play an important role defining the polarity which defines β-cell organization *in situ* (17). Fusion events *in vitro* also appear nonrandom in insulinoma cells and rodent β-cells, where exocytosis may occur at sites of recently fused vesicles (18–22). Indeed, syntaxin1A and SNAP-25 cluster at sites of secretory granule docking and exocytosis in single β-cells (23, 24) and reduced SNAP-25 and syntaxin1A clusters correlate with impaired fusion (25). Also, fusion sites are localized to sites of Ca^2^+ entry in β-cells, and this interaction is disrupted in β-cells from models of T2D (26, 27) and β-cells of human donors with T2D (28). The extent to which fusion events can repetitively occur at defined ‘hotspots’ on the β-cell membrane, the underlying mechanism for this, and its potential dysregulation in human T2D, remains unclear.

Among the ion channels that control β-cell function, the voltage-dependent L-type Ca^2+^ channels increase intracellular Ca^2+^ and trigger exocytosis (29). Repolarization of pancreatic β-cell action potentials is mediated by voltage-dependent K^+^ (Kv) channels such as Kv2.1 (30). Interestingly, Kv2.1 also forms multi-channel clusters at the plasma membrane (31) that directly facilitate efficient insulin exocytosis (19, 32, 33), may also recruit SNARE complex proteins such as syntaxin1A, SNAP-25 and Munc18-1 (34), and serve as points of contact between the plasma membrane and endoplasmic reticulum (35). The channel regulates exocytosis independent of β-cell electrical activity by binding syntaxin1A at its C-terminus (33, 36), a region that is modified by post-translational SUMOylation (37). A glucose-regulated deSUMOylation at the plasma membrane appears particularly important for the regulation of insulin secretion (38), and when stimulated can rescue exocytotic responses in human T2D β-cells (11).

Here we examined exocytotic responses from β-cells of 80 human donors with and without T2D. We demonstrate an impaired insulin secretion that can occur independent of a reduced islet insulin content in this cohort and confirm a single β-cell impairment in exocytotic function. By total internal reflection fluorescence (TIRF) microscopy and spatial analyses, we demonstrate a non-random distribution of fusion events across the cell membrane and show that this patterning is regulated by glucose and is disrupted in β-cells of donors with T2D. Knockdown of Kv2.1 reduces the spatial organization of these events, and we show that channel clustering controls the density of fusion ‘hotspots’ while the SUMOylation status of the channel at C- and N-terminal sites regulates the overall proportion of fusion events that occur at these sites. Finally, up-regulation of Kv2.1 in T2D β-cells can convert a random distribution of fusion events into a compartmentalized pattern similar to that seen in β-cells of donors without diabetes.

## RESULTS

### Impaired spatial patterning of fusion in β-cells from donors with T2D

Glucose-stimulated insulin secretion is impaired from islets of donors with type 2 diabetes (T2D) (6, 39). In a cohort of 80 human islet preparations (**Suppl. Tables 1-2**) we confirm that glucose-stimulated insulin secretion is impaired in T2D and show that this can occur without a reduction in insulin content (**Fig 1A-B**). Although we (40) and others (39) have reported reduced islet insulin content from donors with long-standing T2D, the disease duration in these donors is relatively short (average 6.7 years, including some ‘undiagnosed diabetes’). Donors with longer duration diabetes, or with higher HbA1c, tended to have lower islet insulin content (not shown). In these donors we find that while increasing glucose facilitates exocytosis in β-cells from donors with no diabetes (ND), seen as a rightward shift in the distribution of exocytotic responsiveness of these cells (Fig 1C), this is impaired in T2D (**Fig 1D**) consistent with reduced function at the level of single β-cells (11).

**Figure 1.**
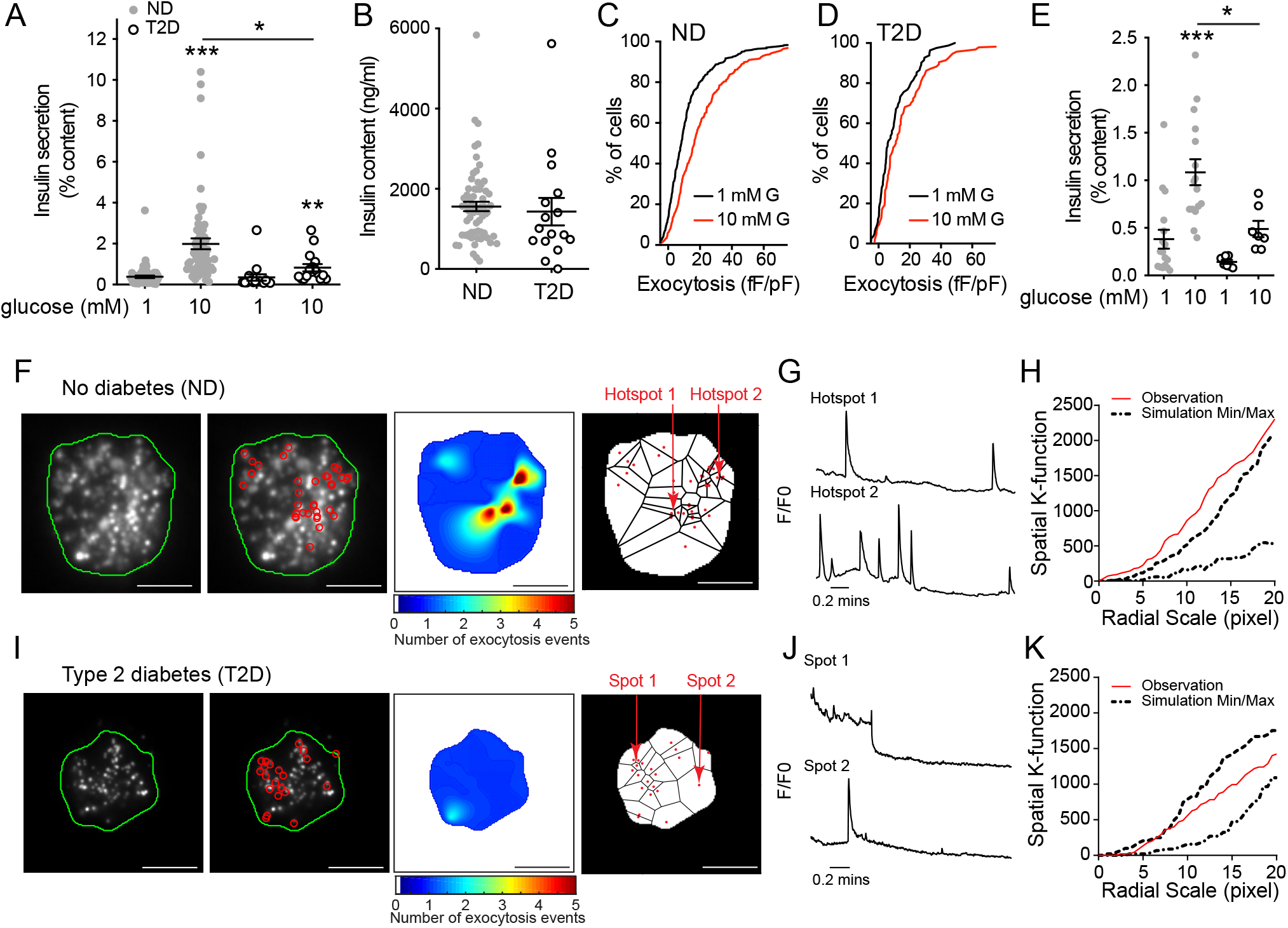
β-cell exocytosis is impaired in T2D. **A)** Compared to islets of donors with no diabetes (ND; n=63), glucose-stimulated insulin secretion is impaired from islets of donors with type 2 diabetes (T2D; n=17). **B)** In these donors (avg. 6.7 years duration) islet insulin content is not different. **C-D)** Cumulative distribution of exocytotic responses of β-cells from the same donors, measured by patch-clamp electrophysiology. Glucose (10 mM; *red*) amplifies the exocytotic responses of ND β-cells (**C**; n=701 cells) but not T2D β-cells (**D**; n=156 cells). **E)** Impaired glucose-stimulated insulin secretion observed in the subgroup of ND (n=17 donors) and T2D (n=7 donors) subsequently used for TIRF imaging. **F)** In individual β-cells expressing NPY-EGFP, fusion events (*red circles*) observed by livecell TIRF microscopy at 5 mM glucose in ND β-cells, a heatmap of fusion event density, and Voronoi diagram used to separate exocytosis sites (scale bar is 5 μm). **G)** Representative recordings from the areas indicated in panel F. **H)** The non-random nature of fusion events in ND β-cells is demonstrated by spatial K-function calculation (*red line*) greater than the simulated (*dashed lines*) maximum. **I-K)** The same as panels F-H, but in a T2D β-cell, where events appear more randomly distributed. Significance was determined by Mann-Whitney test (panel B) or the Kruskal-Wallis one-way analysis of variance followed by Mann-Whitney post-test (panels A, E). *P<0.05; **p<0.01; and ***p<0.001.

Impaired exocytosis in models of T2D is associated with an uncoupling of exocytosis from sites of Ca^2+^ entry (26, 27) and human T2D β-cells have a defect in glucose-dependent granule docking (12). We performed live-cell total internal reflection fluorescence (TIRF) microscopy of single dissociated β-cells to assess the spatial relationship of fusion events in cells from a subset of ND and T2D donors. In this subset, we confirmed impaired insulin secretory function associated with T2D (**Fig 1E**). To identify and perform spatiotemporal analysis of vesicle fusion, a MATLAB-based software (18) was applied to live-cell recordings obtained with over two minutes in a bath solution with 5 mM glucose, a concentration shown to facilitate insulin exocytosis (11), following preincubation at 1 mM glucose for 30 minutes. Fusion events from ND β-cells had a higher degree of spatial organization than those observed in T2D b-cells, with numerous events appearing to occur repeatedly at ‘hotspots’ (**Fig 1F-K; Suppl Movie 1-2**) within a defined area of 4×4 pixels determined by average vesicle diameter (see Methods). Each apparent hotspot was confirmed by generating event heat maps and comparing relative fluorescence to the first frame (**Fig 1G,J**). The spatial organization of events was also confirmed by calculating Ripley’s K-function values, which demonstrate a non-random distribution of these events (**Fig 1H,K**). In ND β-cells 41.6% of events occur within spatially localized regions, while only 18.3% do so in T2D β-cells.

We confirmed impaired exocytosis by patch-clamp electrophysiology at 5 mM glucose (**Fig 2A**). From the same donors, live-cell imaging of β-cells expressing NPY-EGFP to mark secretory granules (**Fig 2B**) at 5 mM glucose (following a 1 mM glucose pre-incubation) reveals that accumulated fusion events are decreased 56.1% in the T2D β-cells (**Fig 2C**). Calculation of a Clustering Index (CI - see Methods) demonstrates that fusion events in the ND β-cells showed significant compartmentalization (‘observation’) compared with Monte Carlo simulation run 100 times of each cell assuming random distribution of events (‘simulation’), or events in the T2D β-cells which were seen to be no more organized than random simulation (**Fig 2D**). While the above analyses account for differences in event frequency, we wished to confirm that the differential organization of events occurs independently of an overall decreased exocytosis since event frequency correlates with the proportion occurring at ‘hotspots’ (**Fig 2E**).

Analysis of a sub-set of cells with equivalent exocytosis event frequency (dashed box and *inset*, **Fig 2E**) demonstrates that the proportion of events occurring at ‘hotspots’ (**Fig 2F**) and the density of these (**Fig 2G**) are decreased in T2D. Thus, impaired insulin secretion from islets of donors with T2D is associated with reduced exocytotic responses measured both by patch-clamp and live-cell imaging, and with an independent reduction in the compartmentalization of fusion events determined by generation of exocytotic heat maps, Ripley’s K-function, Voronoi polygon-derived Clustering Index, and cell sub-set analyses.

**Figure 2.**
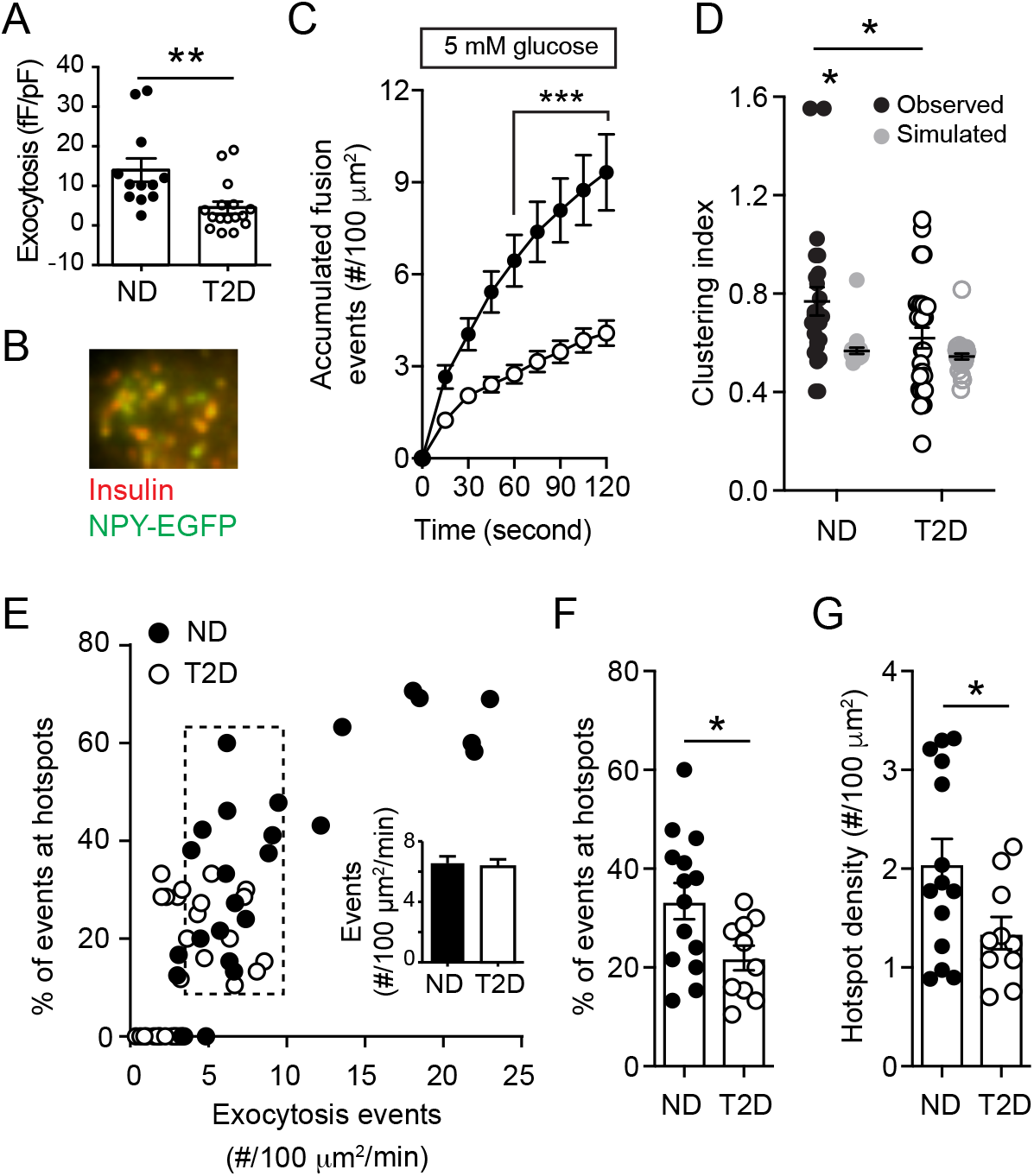
Spatial compartmentalization of granule fusion is reduced in T2D. **A)** Exocytosis measured by patch-clamp in a sub-set of donors at 5 mM glucose, demonstrates impaired function in T2D (n=12 and 16 cells from ND 3 and 3 T2D donors). **B-C)** In □-cells expressing NPY-EGFP, which co-localizes with insulin by immunostaining (M), fusion events were monitored by TIRF microscopy also at 5 mM glucose (ND, *black circles*, n=16 cells from 5 donors; T2D, *open circles*, n=26 cells from 6 donors). **D)** Uniform index of events from panel N calculated (*black*) from Voronoi polygon areas and compared with random simulations (*grey*). **E)** Scatter plot of total fusion events versus the proportion of which occur within clusters. A subset of cells (*dashed box*) with equivalent exocytosis (*inset*) were selected for further analysis. **F)** The proportion of events occurring in spatially defined clusters, and; **G)** The density of these clusters. Significance was determined by the Student’s t-test (panel A,F), by ANOVA and Bonferroni post-test (panel C), by or by the Kruskal–Wallis one-way analysis of variance followed by Mann-Whitney post-test (panels D,G). *P<0.05; **p<0.01; and ***p<0.001.

### Glucose-dependent spatial patterning of exocytosis is impaired in T2D β-cells

In addition to its ability to initiate β-cell action potential firing, glucose signaling amplifies depolarization-induced insulin secretion and β-cell exocytosis (11, 41). Glucose, but not direct depolarization with KCl, can diminish submembrane granule turnover in mouse β-cells (42). We stimulated β-cells with depolarizing KCl at either 1- or 5-mM glucose. Temporal analysis reveals that exocytosis is impaired in T2D β-cells under both conditions (**Fig 3A, B**). At 1 mM glucose, the spatial uniformity of fusion events in ND β-cells was not different than that predicted by random simulation, but the compartmentalization of events increased in the presence of 5 mM glucose (**Fig 3C**). Examining β-cells with equivalent rates of exocytosis in ND and T2D (**Fig 3D**, *dashed box*), KCl-stimulated exocytosis at 5 mM glucose showed an increased proportion of fusion events at ‘hotspots’ (**Fig 3E**), along with an increased density of these sites (**Fig 3F**), effects that are blunted in β-cells of donors with T2D (**Fig 3E-F**).

**Figure 3.**
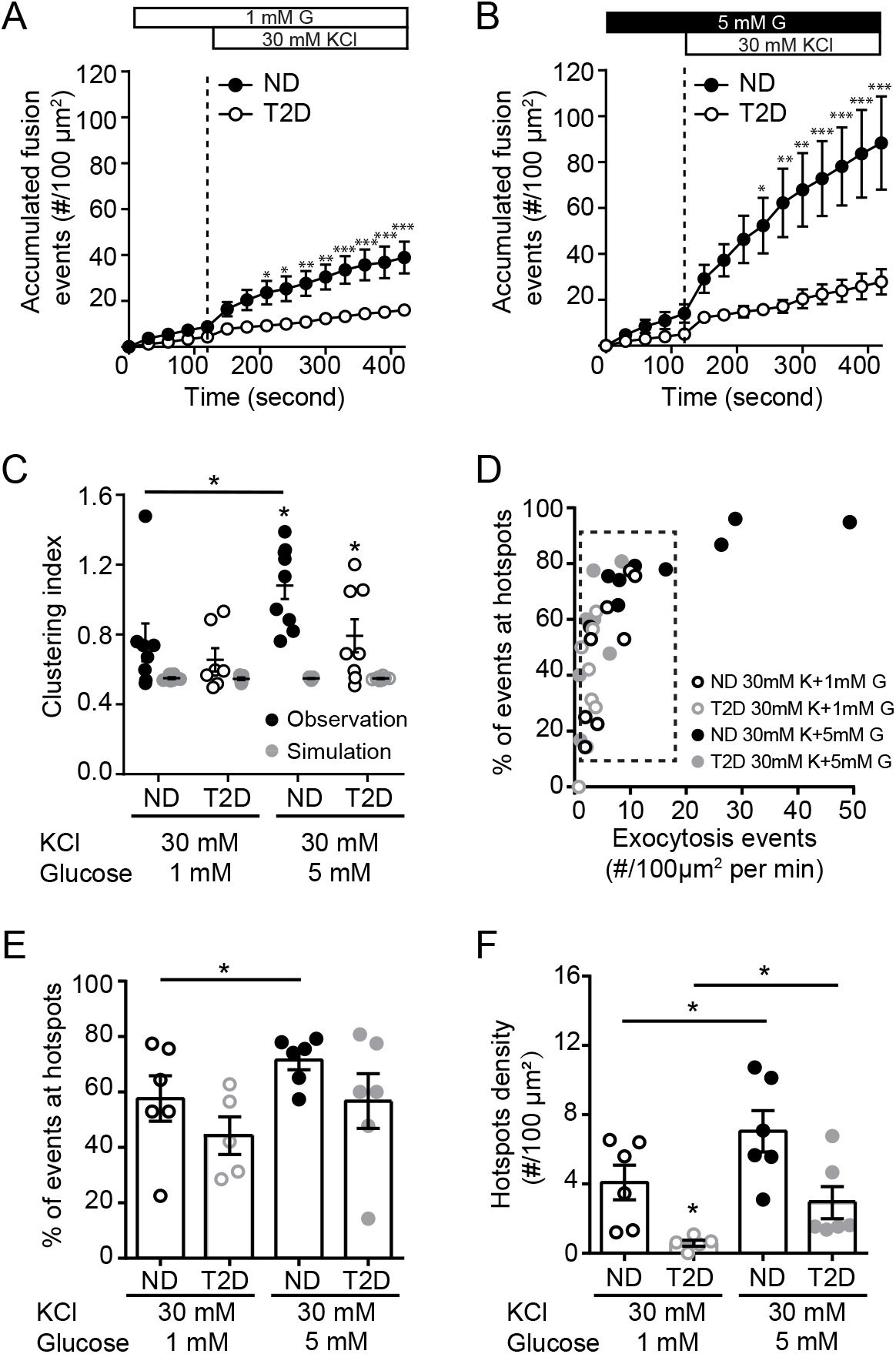
Glucose regulates the spatial organization of fusion events. **A-B)** Accumulated fusion events monitored by TIRF microscopy of ND β-cells (*black circles*) and T2D β-cells (*open circles*) expressing NPY-EGFP, in response to KCl at 1 mM (A; n=7 and 8 cells from 6 and 3 donors) and 5 mM glucose **(B**; n=9 and 9 cells from 6 and 3 donors). **C)** The uniform index is increased at 5 mM glucose in the ND (*closed symbols*), but not T2D (*open symbols*), β-cells. **D)** Scatter plot showing total fusion events versus the proportion of events in spatially restricted clusters. Cells with similar exocytosis (*dashed box*) used for further analysis. **E-F)** The proportion of events occurring in organized clusters (**E**) and the density of these clusters (**F**) is increased at 5 mM glucose in ND β-cells, but impaired in T2D β-cells. Significance was determined by ANOVA and Bonferroni post-test. *p<0.05, **p<0.01, ***p<0.001.

### Observation of fusion occurring near sites of concurrent or recent membrane-resident granules

Insulin granule docking is a critical glucose-dependent step in insulin secretion, and is dysregulated in T2D (12). Considering that glucose-stimulation recruits and KCl-stimulation depletes docking granules (43), together with our observation that glucose increases the proportion and density of ‘hotspots’, we hypothesized that membrane-associated granules may mark sites where repeated exocytosis occurs. Many fusion events throughout our recordings overlap with sites of membrane-resident granules observed at the initiation of imaging (**Fig S1; Suppl Movie 3**). As illustrated in **Fig 4A** (and **Suppl Movie 4**) we observe fusion events that happen both around pre-docked granules and, later in the same recording, at this site after the pre-docked granule had fused. We have excluded events from this analysis that occur within very rapid (<1 s) succession directly overlaid with a docked granule (**Fig 4B**) since this may represent ‘flickering’ or ‘kiss and run’ of an individual granule.

**Figure 4.**
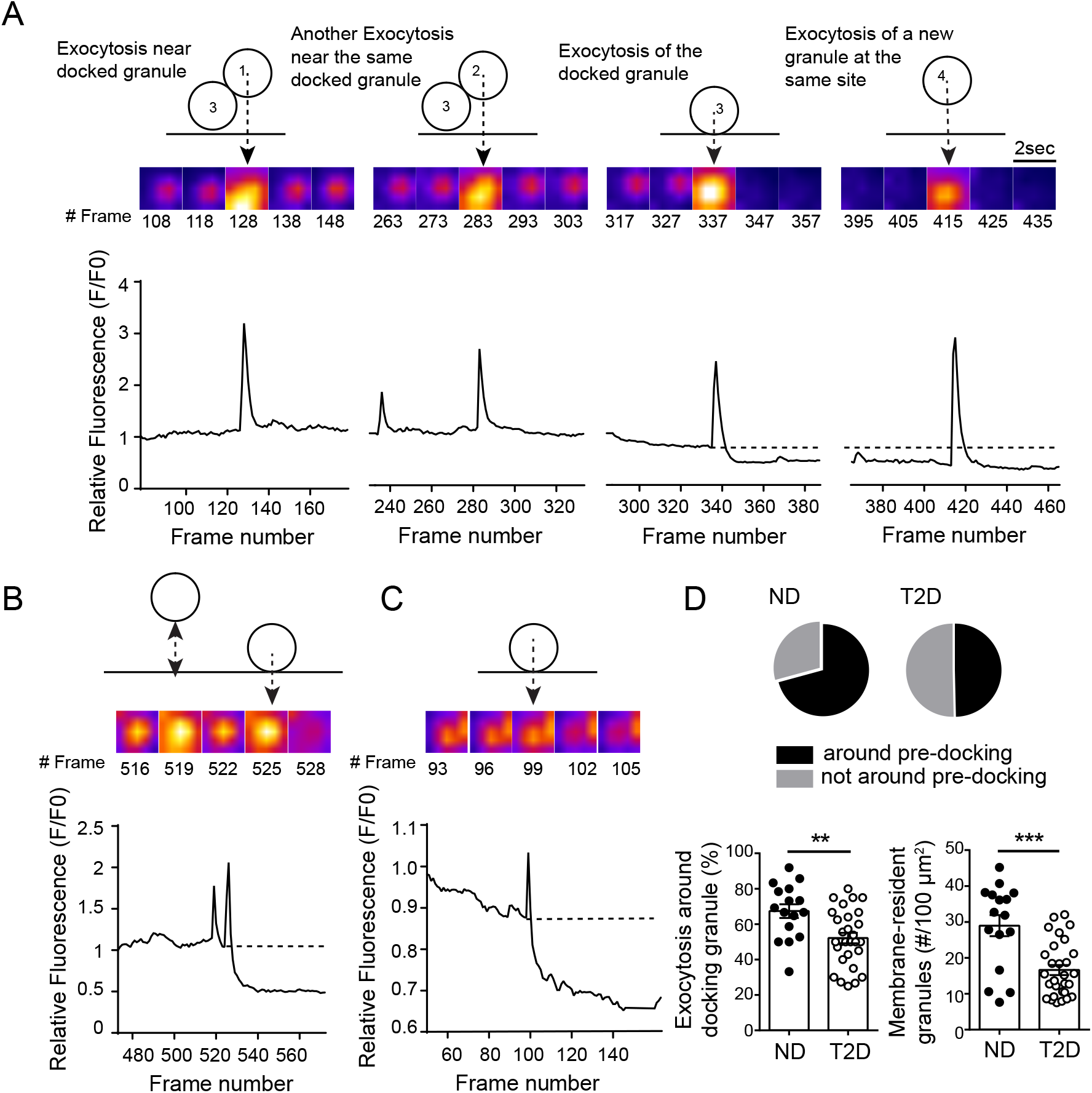
Fusion occurs around sites of concurrent or recent membrane-associated granules. **A)** Exocytosis occurring around a membrane-localized granule in a ND β-cell. Two events occur adjacent to a membrane-resident granule, followed by fusion of that granule itself, and then a new fusion event at the same site. **B)** Rapid flickering events before full fusion, illustrated here, are ignored in this analysis. **C)** In T2D β-cells most fusion occurred as individual, rather than as sequential, events. **D)** Quantification of the proportion of events occurring around initially membrane-localized granules (*pie charts, and bottom left*) and the initial density of membrane-resident granules (*bottom right*; n=16 and 28 cells from 3 ND and 3 T2D donors). Significance was determined by the Student’s t-test. **p<0.01, ***p<0.001.

Sequential fusion events are less often observed in T2D β-cells, which is dominated by isolated events (**Fig 4C**). We find in ND β-cells (5 mM glucose) that 67.4% of fusion events occur at sites marked by membrane-associated granules present within the first frame of the recording, including the fusion of that docked granule itself, while only 55.2% of exocytosis in T2D β-cells occurs near such sites (**Fig 4D**). Importantly, the overall density of membrane-associated granules is decreased 35.8% in T2D β-cells (**Fig 4D**), which is consistent with a previous report (12).

### The density of membrane resident granules, and fusion ‘hotspots’, is controlled by Kv2.1

We demonstrated previously in insulinoma cells that the formation of multi-channel clusters of Kv2.1 controls the density of membrane-associated granules (32) and that fusion events occur on the periphery of Kv2.1 clusters (19). Knockdown of Kv2.1 expression in human β-cells (**Fig 5A**) reduces the density of pre-docked secretory granules (**Fig 5B**) and the frequency of fusion events observed at 5 mM glucose (following a 1 mM glucose preincubation) (Fig 5C) but not the overall proportion of those events that occur near the pre-docked granules (**Fig 5D**). The compartmentalization of fusion events, determined by calculation of Clustering Index, was not significantly reduced following Kv2.1 knockdown (**Fig 5E**). Similar results are observed upon analysis of a subset of cells (*dashed box*, **Fig 5F**) with similar levels of exocytosis (**Fig 5G**), where the proportion of exocytotic events occurring at hotspots was unchanged (**Fig 5H**), but the overall density of these hotspots was decreased (**Fig 5I**).

**Figure 5.**
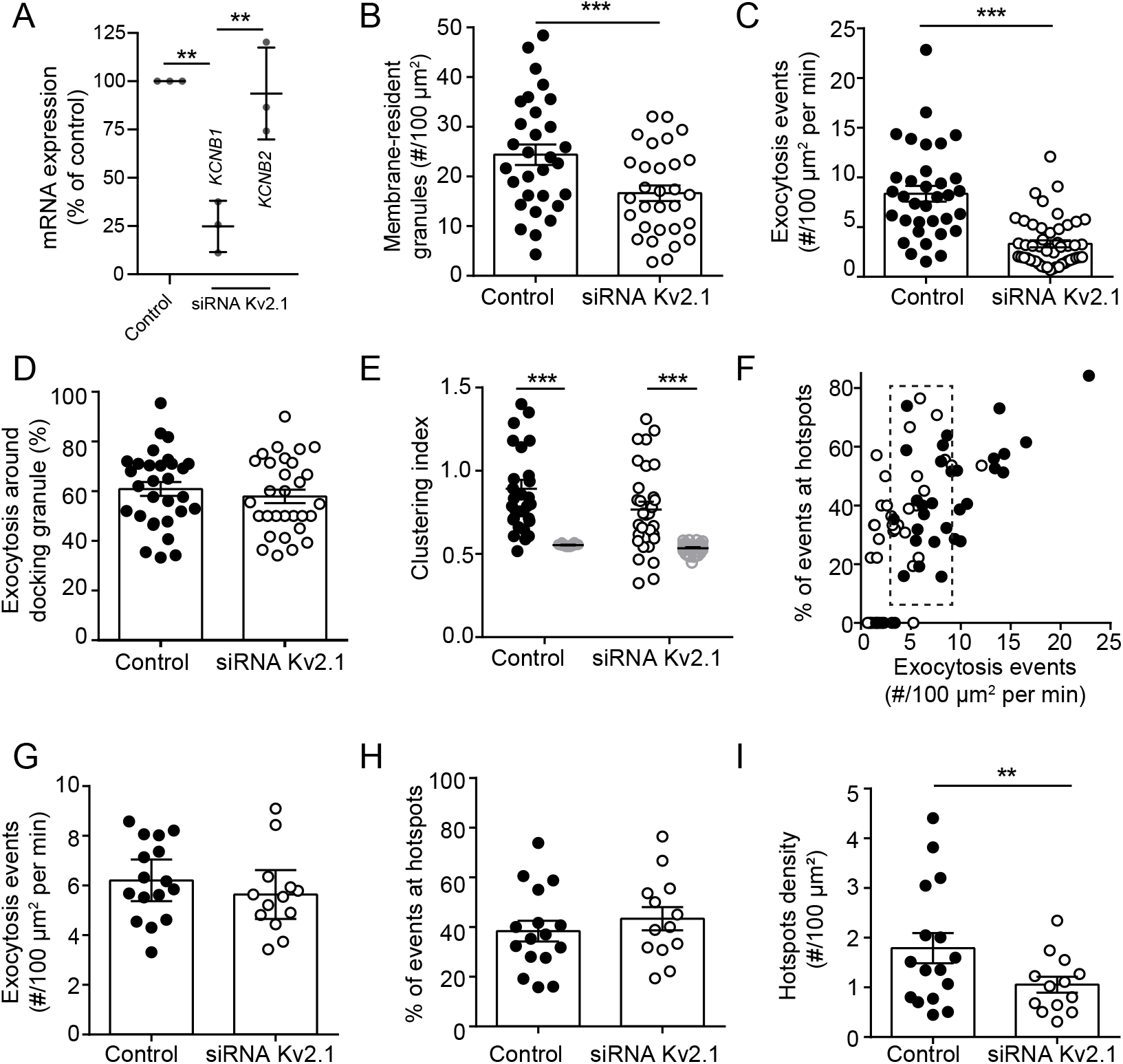
Kv2.1 regulates the density of membrane-resident granules and clustered fusion sites. **A)** Knockdown of Kv2.1 expression by siRNA in human islet cells confirmed by qPCR of Kv2.1 (*KCNB1*) and the related Kv2.2 (*KCNB2*). **B)** The initial density of membrane-resident granules marked by NPY-EGFP is reduced by Kv2.1 knockdown, as is; **C)** the frequency of exocytotic events. **D)** This occurs without a change in the overall proportion of fusion events occurring at sites marked by membrane-resident granules (n=30 and 30 cells from 3 donors). **E)** The spatial organization of these events is only modestly decreased. **F-I**) Examination of a subset of cells (**F**; *dashed box*) with similar event frequency (**G**) demonstrates that the proportion of events occurring in spatially clustered regions is unchanged by Kv2.1 knockdown (**H**) while the density of these sites is decreased (**I**). Significance was determined by ANOVA and Bonferroni post-test (panel A), by the Student’s t-test (panels B-D,G-I), or by the Kruskal–Wallis one-way analysis of variance followed by Mann-Whitney post-test (panel E). *p<0.05, **p<0.01, ***p<0.005.

The formation of multi-channel clusters is required for the facilitation of β-cell exocytosis by Kv2.1 (19, 32). Expression of a C-terminal truncated channel (Kv2.1-∆C318) shown by us and others to be clustering-deficient (32, 44) in ND β-cells reduced the compartmentalization of fusion events (**Fig 6A**). In a subset of ND cells with equivalent exocytosis (**Fig 6B**, *dashed box*), up-regulation of wild-type Kv2.1 did not effect the spatial organization of fusion events, while the Kv2.1-∆C318 decreases proportion of events occurring at hotspots in ND β-cells (**Fig 6C**) without affecting the overall hotspot density compared with cells expressing mCherry alone (**Fig 6D**). In T2D β-cells, up-regulation of the wild-type channel restored the spatial compartmentalization of events (**Fig 6E-H**). We find that up-regulation of ‘clustering-sufficient’ wild-type Kv2.1 increases membrane-resident secretory granules by 30.0%, and increases exocytosis at these sites by 42.4% compared to the mCherry control group in ND donors (**Fig S2A**). Similarly, in T2D donors the wild-type Kv2.1, but not the ‘clustering deficient’ Kv2.1-ΔC318, can increase membrane-associated granules by 37.5% and increase fusion events around these by 50%(**Fig S2B**).

**Figure 6.**
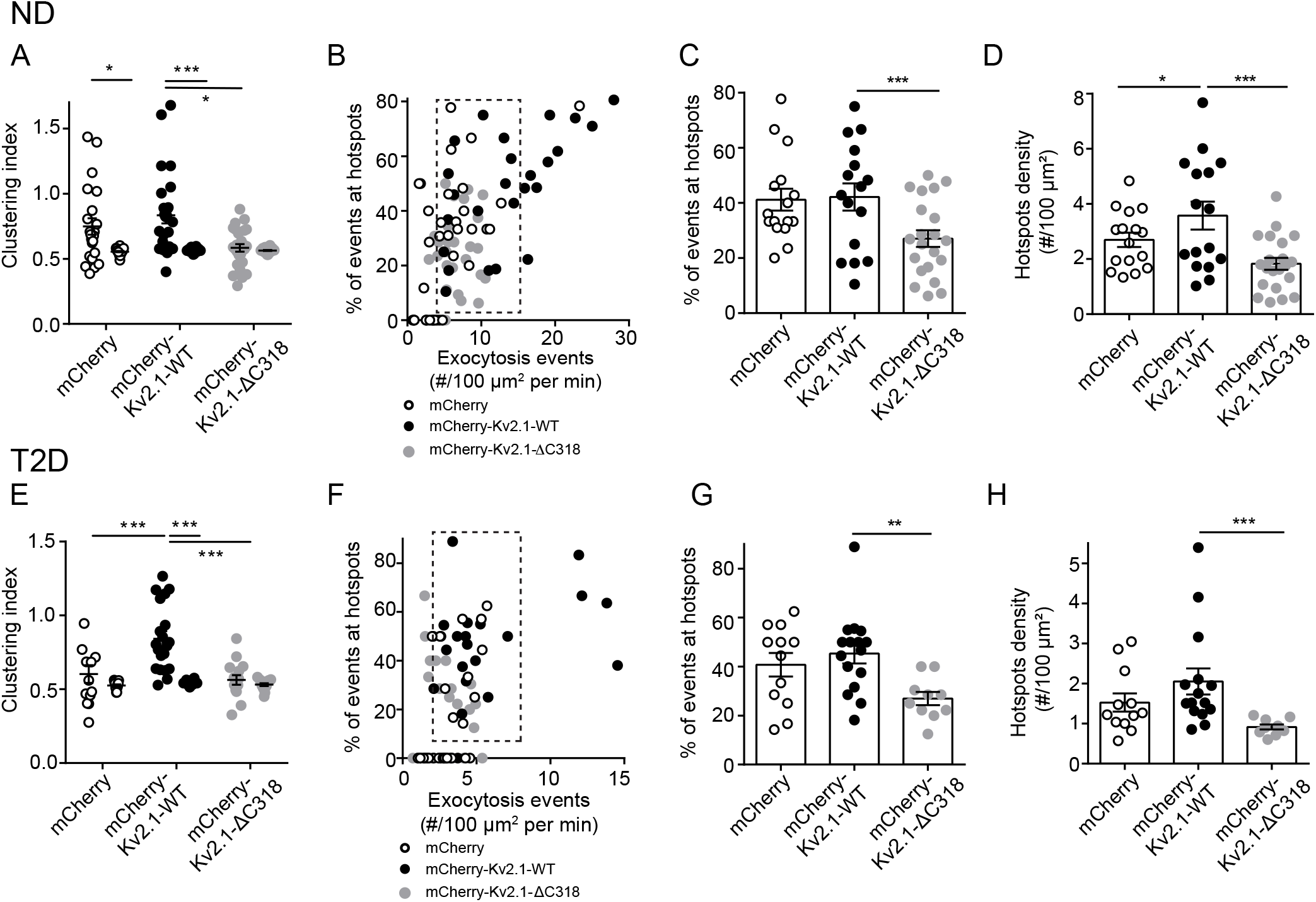
Upregulation of Kv2.1 restores the spatial organization of fusion sites in T2D β-cells. **A)** In ND β-cells, compared with the full-length channel (Kv2.1-WT) a clustering deficient mutant (Kv2.1-∆C318) reduced the uniformity of fusion events monitored by TIRF imaging of granule-targeted NPY-EGFP (n=22, 26 and 24 cells from 3 donors). Cells expressing the channel constructs were identified by channel-tagged mCherry and compared to cells expressing mCherry alone. **B-D)** In a subset of cells with equivalent rates of exocytosis (**B**, *dashed box*) expression of Kv2.1-WT had little effect on the proportion of events within hotspots (**C**) and slightly increased their density (**D**), while Kv2.1-∆C318 reduced the overall contribution of hotspots to exocytosis. **E-H)** Same as panels A-D, but in T2D β-cells (n=12, 16 and 10 cells from 3 donors). Here, Kv2.1-WT rescued the compartmentalization of fusion events. In a sub-set of cells with similar fusion frequency (**F**; *dashed box*) the overall proportion of events occurring in these spatially restricted regions (**G**) and the density of these sites (**H**) are unchanged by WT-Kv2.1. Significance was determined by the Kruskal–Wallis one-way analysis of variance followed by Mann-Whitney posttest (panels A,H), or by ANOVA and Bonferroni post-test (panels C-E,G). *p<0.05, **p<0.01 and ***p<0.001.

### SUMOylation sites on Kv2.1 determine the proportion of fusion events occurring in hotspots

The clustering of Kv2.1 contributes to the spatial coordination of fusion events secondary to the control of membrane-resident granule density. We (45) and others (37) have shown that Kv2.1 is regulated by post-translational SUMOylation, which has also been shown to negatively regulate insulin exocytosis (38). The most well established SUMOylation site on the channel is located at lysine 470 (37) within a syntaxin 1A binding domain of the C-terminus (46) previously shown to have a role in β-cell exocytosis (33). A second SUMOylation motif, at lysine 145 within a potential SNAP-25 binding domain in the N-terminus (47), is more highly conserved across species (**Fig S3A**) and homologous to a demonstrated SUMOylation site in other Kv channels, such as Kv1.1, Kv1.2 and Kv1.5 (48) (**Fig S3B**). Mutation of K145 does not appear to alter Kv2.1-mediated K^+^ currents (37), which we confirm (**Fig 7A,B**). However, loss of the K145 SUMOylation site (Kv2.1-K145R) blunts the suppressive effect of SUMO1 on exocytosis in insulinoma cells (**Fig 7C-F**) and when combined with abolishing the K470 SUMOylation site (Kv2.1-K145/470R) completely prevents SUMO1-dependent suppression of exocytosis (**Fig 7F**).

**Figure 7.**
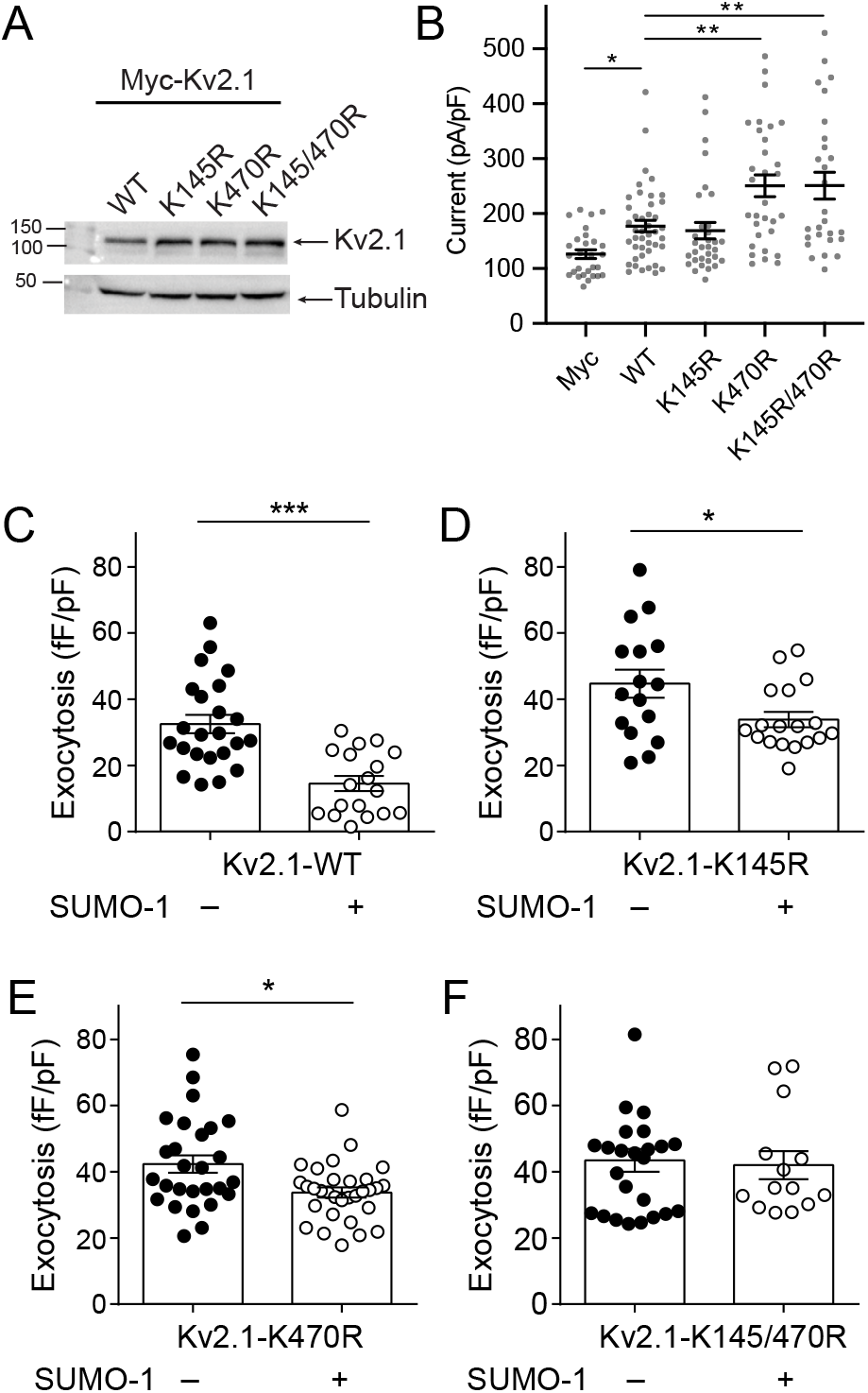
SUMOylation sites on the Kv2.1 N- and C-termini control β-cell exocytosis. **A)** When expressed in HEK293 cells mutation of Kv2.1 C-terminal (K470R) or the N-terminal (K145R) SUMOylation sites, alone or in combination, has no effect on channel protein level (representative of 3 experiments). **B)** When expressed in INS 832/13 cells loss of the K145 site has no effect on channel current measured by whole-cell patch-clamp electrophysiology, while currents are increased upon mutation of K470 or of both sites in concert (n=16, 20, 27, 20, 28 cells). **C-F**) When examined by measurement of cell capacitance responses, SUMO1 suppresses exocytosis in INS 832/13 cells expressing Kv2.1-WT (**C**; n=23, 18 cells), and this effect is reduced in cells expressing Kv2.1-K145R (**D**; n=16, 18 cells) or Kv2.1-K470R (E; n=27, 29 cells). Mutation of both SUMOylation sites (**F;** Kv2.1-K145/470R; n=24, 14 cells) abolished the inhibitory effect of SUMO1 on exocytosis. Significance was determined by the Kruskal–Wallis one-way analysis of variance followed by Mann-Whitney post-test (panels B), by the non-parametric Mann-Whitney test alone (panels C,E), or by the Student’s t-test (panels D,F).*p<0.05, **p<0.01 and ***p<0.001.

In human β-cells from donors without diabetes, increasing SUMO1 reduces the compartmentalization of fusion events (**Fig 8A,B**). Loss of Kv2.1 SUMOylation sites in the Kv2.1-K145/470R double mutant prevents this (**Fig 8C,D**). We find that SUMO1 reduces the compartmentalization of events such that the Clustering Index is no longer different from random simulations and this is completely prevented in the Kv2.1-K145/470R group (**Fig 8E**). Finally, in cells with equivalent rates of exocytosis (**Fig 8F**, *dashed box*), SUMOylation of Kv2.1 reduces the proportion of fusion events occurring within hotspots (**Fig 8G**) with only minimal impact on total hotspot density (**Fig 8H**). Altogether, we find that multi-channel Kv2.1 clusters promote the recruitment of membrane-resident secretory granules which define sites of compartmentalized fusion. The actual targeting of granules to these sites depends on deSUMOylation of the channel (**Fig S4**).

**Figure 8.**
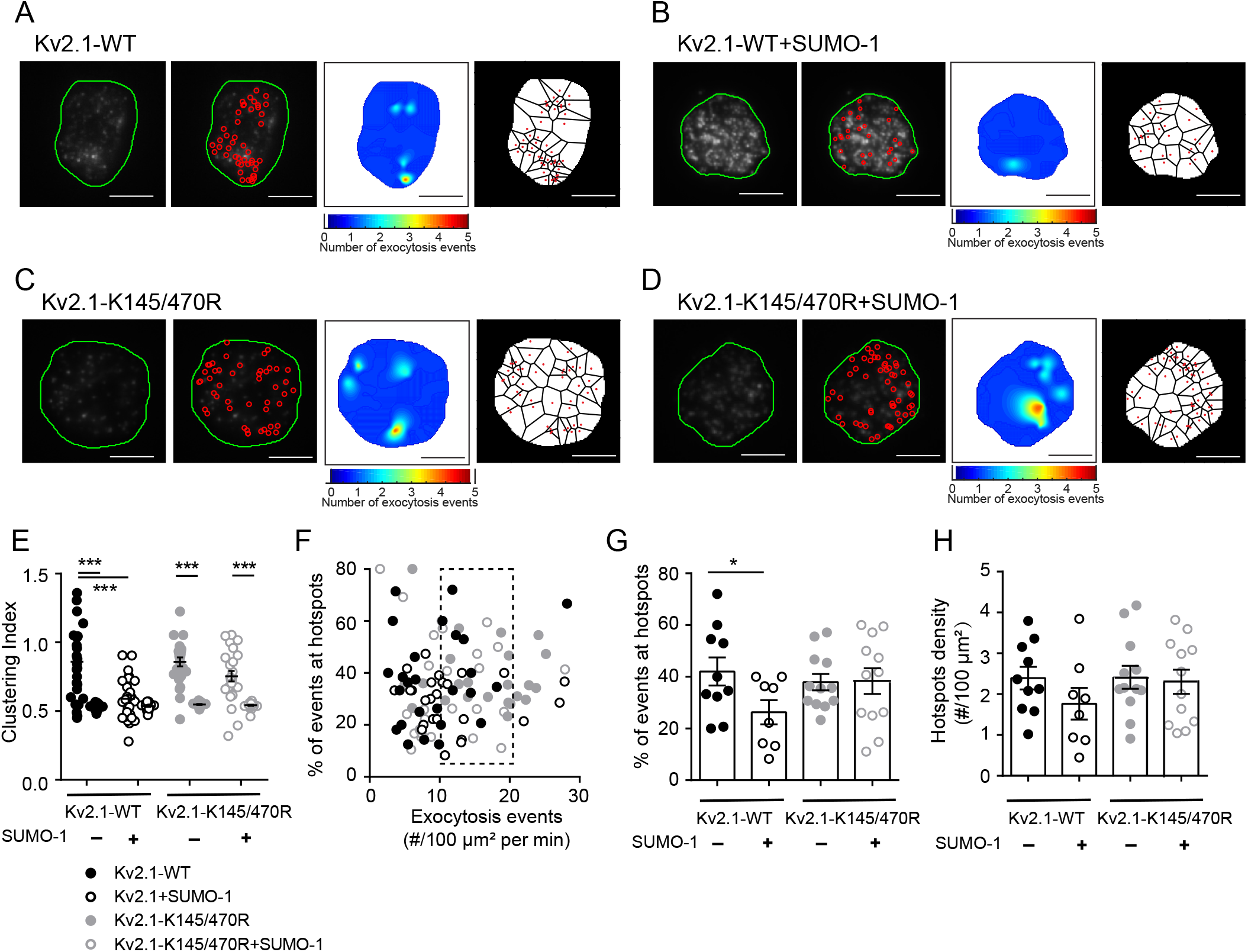
Kv2.1 SUMOylation sites regulate compartmentalization of fusion events. **A-D)** Example fusion events tracked by TIRF imaging of NPY-EGFP expression shown individually (*red circles*), as density heatmaps, or separated by Voryoni polygons in ND β-cells expressing Kv2.1-WT alone (**A**) or with SUMO1 (**B**), and Kv2.1-K145/470R alone (**C**) or with SUMO1 (**D**). Scale bars are 5 μm. **E)** Uniformity index values calculated from Voryoni polygons show that SUMO1 reduces the compartmentalization of fusion events in cells expressing Kv2.1-WT, but not the SUMOylation-deficient mutant Kv2.1-K145/470R. (n=10, 8, 12, 12 cells from 4 donors) **F)** Scatterplot showing the relationship between proportion of events occurring within hotspots versus event frequency. A subset of cells (*dashed box*) was chosen for further analysis. **G-H**) SUMO1 reduced the proportion of events occurring at hotspots (**G**) with little impact on the overall hotspot density (**H**). Significance was determined by ANOVA followed by Bonferroni post-test.*p<0.05, **p<0.01 and ***p<0.001.

## DISCUSSION

The regulated exocytosis of insulin from human pancreatic β-cells is impaired in T2D. Indeed, reduced exocytotic function is observed even in β-cells of non-diabetic human donors with genetic risk for T2D (10). Several factors likely contribute to this lower exocytotic responsiveness, including reduced SNARE expression (8), a loss of regulatory metabolic signals (11), impaired microtubule (49) or actin (50) regulation of granule access to the membrane, and the uncoupling of granule fusion from sites of Ca^2+^ entry. The latter has been described in models of free fatty acid culture (26), high fat fed mice (27), and in β-cells from human T2D donors (28). Recently, the glucose-dependent recruitment of granules to docking sites was shown to be a limiting step in insulin secretion which is disturbed in T2D (12). The spatial organization of events at the plasma membrane, such as the localization of integrin signals (16) or the clustering of Ca^2+^ channels (28), are important determinants of regulated insulin secretion. However, little is understood about the control of the spatial patterning of fusion events in human health and T2D.

An insulin secretory deficit, including a reduced responsiveness to glucose, is an important determinant of T2D development and progression (51). While impaired islet mass may contribute to reduced insulin secretion in T2D, at least later in the disease (5), the contribution of reduced islet insulin content versus impaired secretory responses early in T2D progression is less well recognized. Although insulin content has been shown to be decreased in T2D human islets by us (40) and others (39), in the present data-set we observed impaired insulin secretory responses without an obvious decrease in insulin content. This is likely due to the relatively short duration of T2D in this cohort (6.7 +/-1.9 years) where only 5/17 donors had reported disease duration of 10 years or more. Of the T2D donors with no information on disease duration, one appeared well-controlled (HbA1c 6.2) and the other was managed with diet only. One additional diet-controlled T2D donor (R198) diagnosed 10 years prior may be considered in remission based on available information (see Methods). Although this donor-to-donor heterogeneity could complicate the interpretation of secretion results, which is the case for all human islet studies, in this paper we have taken a step forward in addressing this by linking each preparation used to its full donor, quality control, and islet function data via a web tool (www.isletcore.ca) and hyperlinks provided in **Supplementary Tables 1 and 2.**

Single-cell impairments in insulin exocytosis from human T2D b-cells (11, 32) may result from reduced glucose-dependent granule docking (12) and disrupted localization of granule fusion to sites of Ca^2+^ entry (26–28). Previous studies have suggested both directional control of insulin release towards the vasculature in situ (13–15) and the existence of repetitive release sites (18–22). Focal adhesion activation may contribute to this (52), resulting from ECM-dependent interactions that drive cell polarity *in situ* (17). While it seems likely that the additional signals such as integrin activation by the ECM-interactions (16) would provide an additional layer of organization, we observe a glucose-dependent spatial patterning of fusion events across the surface of isolated β-cells that are defined by the presence of previously or concurrently docked granules. The observation that membrane-docked granules mark ‘hotspots’ in β-cells suggests that the reduced organization of fusion events in T2D β-cells could be secondary to impaired glucose-dependent granule docking. In T2D donors reduced glucose-dependent docking is suggested as a major upstream defect (12), and we also report a reduced density of membrane-associated granules in T2D β-cells.

SNARE proteins and ion channels are compartmentalized across the cell membrane, in complexes which are generally thought to facilitate efficient exocytosis. Previously, we reported that Kv2.1 in pancreatic β-cells co-localizes with docking granules (19, 32) and directly facilitates exocytosis independent of its electrical function (36) in a manner that is dependent upon the formation of multi-channel clusters (32). Indeed, we recently demonstrated that fusion events in insulinoma cells occur adjacent to Kv2.1 clusters (19) and suggested that these sites act as ‘reservoirs’ for the provision of release-competent granules. This role is supported by our observation that depletion or over-expression of Kv2.1 reduces or increases, respectively, the density of membrane-resident granules and consequently the density of fusion hotspots (**Fig S4**). These manipulations have little or no effect on the overall proportion of total fusion events occurring within these hotspots, suggesting that while clustered Kv2.1 channels contribute to setting the number of hotspots another mechanism is involved in directing fusion to these sites.

SUMOylation is an important regulatory mechanism for membrane proteins. Kv2.1 can by SUMOylated (37, 45), and we find an important functional effect of manipulating SUMOylation sites in both the N- and C-terminus of the channel. The N-terminal SUMOylation site is less well established and has no apparent impact on the biophysical function of K+ conductance, in line with a previous report (37). We do however find that this is more highly conserved between species and amongst Kv isoforms. This site in Kv1.5 has a demonstrated function in SUMO-control of the channel (48) and here we show that mutation of the consensus lysine residue (K145) reduces the inhibitory effect of SUMOylation on exocytosis in concert with the C-terminal site (K470). This has only minimal impact on the density of hotspots, but instead controls the occurrence of fusion at these sites (**Fig S4**). It seems that Kv2.1 SUMOylation, which occurs at sites of syntaxin-1A/-3 (53) and SNAP-25 (47) binding might not alter the ability for granules to be recruited to the ‘reservoir’, but instead impacts the transfer of granules to the fusion sites. This, and the roles in sequential insulin granule fusion for Munc18b (20) and syntaxin-3 (54), which bind the channel (19), suggest that the likely mechanism for the sequential fusion at hotspots involves Kv2.1 interactions with an alternate SNARE complex.

In summary, we demonstrate a glucose-dependent spatiotemporal compartmentalization of insulin granule fusion events in human β-cells. This contributes to the amplification of insulin exocytosis by glucose and is perturbed in T2D. The voltage-gated K+ channel Kv2.1 plays a structural role in this process, possibly as a component of a complex serving as a ‘reservoir’ of granules for future fusion. Multi-channel clusters of Kv2.1 sets the density of these ‘hotspot platforms’, while deSUMOylation of the channel determines the progression of insulin granules to fusion-competence at these sites.

## METHODS

### Human islets and cell culture

Human islets were isolated at the Alberta Diabetes Institute IsletCore according to procedures deposited in the protocols.io repository (55). A total of 63 donors with no-diabetes (ND; **Suppl Table 1**) and 17 donors with type 2 diabetes (T2D; **Suppl Table 2**) were examined in this study. Extended donor, organ processing, and quality control information is available via hyperlinks for each donor in the Supplementary Tables or by searching relevant donor numbers at www.isletcore.ca. The presence of T2D was determined by clinical reporting at the time of organ procurement, or an HbA1c >6.5% measured from blood tubes provided with the shipped organ. Prior to experiments, islets were cultured in low-glucose (5.5 mM) DMEM with L-glutamine, 110 mg/l sodium pyruvate, 10% FBS, and 100 U/ml penicillin/streptomycin. One donor (R198), with a prior T2D diagnosis, appears to have been in remission with diet-based management but was retained in the T2D group since islet phenotyping demonstrated impaired insulin secretion and β-cell exocytosis.

Human embryonic kidney (HEK) 293 cells from ATCC were cultured in DMEM with 20 mM glucose, 10% FBS, 100 units/mL penicillin, and 100 mg/mL streptomycin at 37°C and 5% CO2. The glucose responsive INS 832/13 insulinoma cell line (56) from Christopher Newgard (Duke University) was cultured in RPMI-1640 with 11.1 mM glucose, 10% FBS, 10 mM HEPES, 0.29 mg/ml L-glutamine, 1 mM sodium pyruvate, 50 μM 2-mercaptoethanol, 100 U/ ml penicillin, and 100 mg/mL streptomycin.

### Adenoviruses, constructs, and treatments

Human pancreatic islets were dissociated using Cell Dissociation Buffer enzyme-free, Hanks’ Balanced Salt Solution (ThermoFisher Scientific. Burlington, ON). Isolated human pancreatic β-cells were infected with adenovirus expressing NPY-EGFP and further cultured for 24-36 h before imaging. We confirmed the localization of NPY-EGFP to insulin granules by immunostaining. Adeno-NPY-eGFP infected human pancreatic β-cells were transfected for 36-48 h with the expression plasmids as outlined for each experiment using Lipofectamine 2000 (Life Technologies, Burlington, ON).

The cDNA encoding wild type rat Kv2.1 or the truncated Kv2.1-∆C318 (Kv2.1 Glu536_Ile853 del) was amplified by PCR using a pcDNA3.1-Kv2.1 plasmid (45) as a template, and inserted between BsrG-I and Xho-I site of Cherry-LacRep plasmid (from Mirek Dundr: Addgene plasmid #18985) by Gibson Assembly to make mCherry-Kv2.1-WT and mCherry-Kv2.1-∆C318.

To generate Myc-tagged Kv2.1 plasmids, the cDNA encoding 5xMyc was inserted between Nhe-I and BsrG-I sites of the mCherry-Kv2.1-WT and mCherry-Kv2.1-∆C318 expression vector by Gibson Assembly. To generate the mutation of SUMOylation sites in Kv2.1, site-directed mutagenesis was performed in a Myc-tagged Kv2.1-WT expression vector to introduce the mutation from lysine to arginine at the position K145, K470 and K145/470. All mutations were confirmed by Sanger DNA sequencing and expression was confirmed by western blotting with anti-Myc primary antibody (1:2000 dilution; Merck Millipore #06-340, Billerica, MA).

The full-size human SUMO1 cDNA, IRES DNA and mCherry cDNA were prepared by PCR using pEYFP SUMO-1 plasmid (from Mary Dasso: Addgene plasmid #13380), pWPI plasmid (from Didier Trono: Addgene plasmid #12254) or Cherry-LacRep plasmid as template, respectively. The pCMV-hSUMO1-IRES-mCherry plasmid was constructed by inserting the cDNA/DNA of Human SUMO1, IRES and mCherry, in this sequence, between Nhe-I and BamH-I sites of Cherry-LacRep plasmid by Gibson assembly. Negative control plasmid, pIRES-mCherry, was prepared in the same manner but without Human SUMO1 cDNA.

Finally, knockdown of KCNB1 expression in human cells was carried out using a mixture of 4 siRNA duplexes (Qiagen; Cat#: S100070777, S100070791, S100070798, S103065930) in which each recognizes different regions of the human KCNB1 gene. Knockdown of KCNB1 was confirmed by qPCR using Taqman expression assays (Assay ID Hs00270657_m1 for KCNB1, Hs00191116_m1 for KCNB2, Applied Biosystems/Thermo Fisher Scientific, MA USA).

### Insulin secretion and electrophysiology

The insulin secretion measurements are detailed in the protocol deposited to protocols.io (57). Glucose concentrations were 1 and 10 mM, after which insulin content was extracted with acid/ethanol. The samples were stored at −20°C and assayed by chemiluminescent immunoassay (STELLUX, Alpco Diagnostics) or electro-chemiluminescent immunoassay (Huaman Insulin Kit, Meso Scale Diagnostics). Experiments were performed in triplicate, and the averaged values were used for analyses. Patch-clamp measurement of Kv currents and exocytotic responses in single HEK293, INS 832/13 or human β-cells, the latter identified by positive insulin immunostaining following the experiment, were performed at 32-35°C as described (33) using a HEKA EPC10 amplifier and PatchMaster Software (HEKA, Lambrecht, Germany).

### TIRFM Imaging

All TIRF imaging used a Cell-TIRF motorized system (IX83P2ZF, Olympus Canada) with a 100x/1.49 NA TIRFM objective, a Photometrics Evolve 512 camera (Photometrics), and Metamorph Imaging software (Molecular Devices). Excitation was at 491 nm (LAS-491-50) and 561 nm (LAS-561-50, Olympus, Germany) with a quad filter passing through a major dichroic and band pass filter (405/488/561/640, Chroma Technology, Bellows Falls, VT). Penetration depth was set to 105 nm, calculated using existing angle of the laser and assuming a refractive index of 1.37. Emission was collected through bandpass filters of 525/25 nm and 605/26 nm for excitations of 488 and 561 nm, respectively. Images were acquired sequentially with single laser excitation to minimize potential bleed-through.

Single β-cells from dispersed human islets were cultured on 25mm round coverslips from Electron Microscopy Sciences (cat# 72225-01) overnight in the human islet media describe above. Live-cell acquisition was 5-Hz with a 200 ms exposure at 35°C. Before acquisition, cells were preincubated (30 mins) in bath containing (in mM) 138 NaCl, 5.6 KCl, 1.2 MgCl_2_, 2.6 CaCl_2_, 5 NaHCO_3_, 1 glucose and 5 HEPES (pH 7.4 with NaOH) and then exposed to 5 mM glucose upon recording. In glucose-stimulation experiments the buffer contained 16.7 mM glucose. For KCl stimulation, the buffer instead contained 30 mM KCl which replaced an equimolar amount of NaCl. Fusion events, indicated by abrupt brightening (ratio of peak fluorescence to background >1.3) and then disappearance of NPY-EGFP fluorescence, were selected and analyzed with a compiled MATLAB (MathWorks) based analysis software (18) within the membrane area (the authors kindly provided their software). Docked granules were detected using a “local maxima” or a weighted centroid method (within a range of 4 × 4 pixels, 0.6328 × 0.6328 μm) (18). Exocytosis events occurring around docking granule were manually selected following overlay of fusion events with identified sites of membrane-resident granules at the start of recordings.

Spatial analysis was performed in MATLAB, 2018a (MathWorks). Voronoi polygons were used to separate each fusion event. The standard deviation of resulting areas divided by the area mean is indicative of the degree of spatial heterogeneity of fusion events. We refer to this, described in Yuan et al. as a ‘uniform index’ (18), as a Clustering Index since increases represent greater compartmentalization of fusion events. These observed values are compared against random simulations run 100 times with the same number of fusion events and the same cell boundaries. Exocytosis hotspots, recognized by MATLAB, was run with a threshold value of 0.6328 μm, which is <4 pixels and represents the minimum resolvable distance to distinguish two granule fusion sites in our system. The ‘% of events at hotspots’ was calculated as the number of fusion events occurring at hotspots divided by the total fusion events and multiplied by 100. The ‘hotspot density (#/100 μm^2^) was calculated as the number of hotspots normalized to cell footprint area. Hotspots visualization heat maps were generated through interpolation (MATLAB ‘v4’ interpolation method) using number of events as the third dimension on a 1-pixel resolution grid.

### Statistical analysis

Data analysis was performed using Fit Master (HEKA Electronik), Origin Lab (v7.0) and GraphPad Prism (v6.0c). All data are shown as the mean ± SEM. Statistical outliers were identified and removed by an unbiased ROUT (robust regression followed by outlier identification) test. Normally distributed data were analyzed by the 2-tailed Student’s t test (for two groups), or ANOVA and Bonferroni post-test (for multiple groups). Data that failed normality tests were analyzed by the non-parametric Mann-Whitney test (for two groups), or the Kruskal–Wallis one-way analysis of variance followed by Mann-Whitney post-test (for multiple groups). A p-value less than 0.05 was considered significant.

### Study approval

All human islet studies were approved by the Human Research Ethics Board (Pro00013094; Pro00001754) at the University of Alberta (Edmonton, Canada) and all families of organ donors provided written informed consent.

## Acknowledgments

We wish to thank Dr. Yongdeng Zhang (Yale University) for providing the computer-assisted analysis software and advice on data analysis. The authors thank the Human Organ Procurement and Exchange (HOPE) and Trillium Gift of Life Network (TGLN) programs for their efforts in procuring pancreas for research. The authors also thank Mr. James Lyon and Mrs. Nancy Smith (Alberta Diabetes Institute IsletCore, University of Alberta) for work isolating human research islets. Human research islet isolations were supported by the Alberta Diabetes Foundation and the University of Alberta. Research was funded by a Foundation Grant from the Canadian Institutes of Health Research (CIHR: 148451) to PEM.

## Author Contributions

JF, JMG, XD, GP, KS, AFS, AB, RJK, DGA, and JEMF researched data; JF, HYG and PEM conceived the study; JF and PEM wrote the manuscript; All authors edited and approved this version.

