## Supplemental Figures for "A glucose-dependent spatial patterning of exocytosis in human β-cells is disrupted in type 2 diabetes"

#### Supplementary Figure Legends

***Figure S1. Fusion events occur at sites of previous or concurrent membrane-resident granules.***

**A)** Membrane-associated granules labelled with NPY-EGFP observed at the beginning of the recording are marked in green (*left*); all subsequent fusion events are shown in red (*center*); fusion events occurring at sites were membrane-localized granules were observed are shown in yellow (*right*). The latter may represent fusion of the membrane-localized granule itself (single yellow event), or of several events at a clustered site (multiple yellow events). Scale bars are 5 μm.

***Figure S2. Upregulation of Kv2.1-WT increases membrane resident granules in ND and T2D.***

**A-B)** Expression of Kv2.1-WT but not Kv2.1-ΔC318 in ND β-cells (**A**) or T2D β-cells (**B**) increases the density of membrane-resident granules and the proportion of fusion events at sites marked by these. Significance was determined by ANOVA and Bonferroni post-test.*p<0.05, **p<0.01 and ***p<0.001.

***Figure S3. Conserved SUMOylation motifs in the N- and C-termini of the Kv2.1 channel.***

**A)** Two previously demonstrated Kv2.1 SUMOylation motifs are conserved across species and between some members of the voltage-dependent $K^+$ channel family. **B)** Overlay of N-terminal domains of Kv2.1 and Kv1.5 showing location of demonstrated and predicted SUMOylation sites.

***Figure S4. The impact of Kv2.1 channel manipulation on fusion event density and spatial organization.***

While Kv2.1 knockdown reduces the overall frequency and density of fusion events in human β-cells, the effects of channel up-regulation depend its SUMOylation status and its ability to form multi-channel clusters. A truncated channel (Kv2.1-ΔC318) that, while electrically functional, cannot form multi-channel clusters and does not increase membrane resident granule and fusion hotspot density (it may even act as a dominant-negative in some measures). SUMOylated Kv2.1 is sufficient to increase membrane granule density but these do not undergo fusion, resulting in a decreased targeting of granules to fusion hotspots.

### Figure. S1

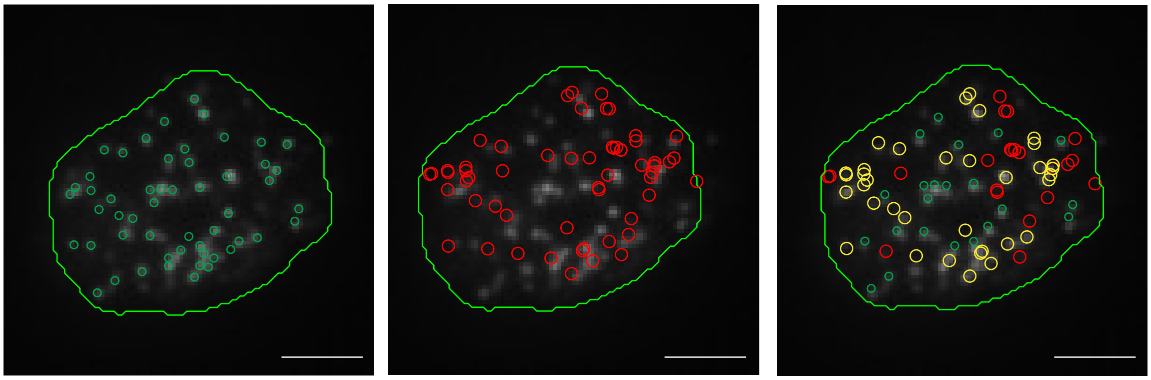

○ Membrane-resident granules marked in first frame

○ Subsequent fusion events

○ Membrane-resident granules that don't fuse
○ Fusion at 'docked granule' sites
○ Fusion at random sites

**Figure. S2**

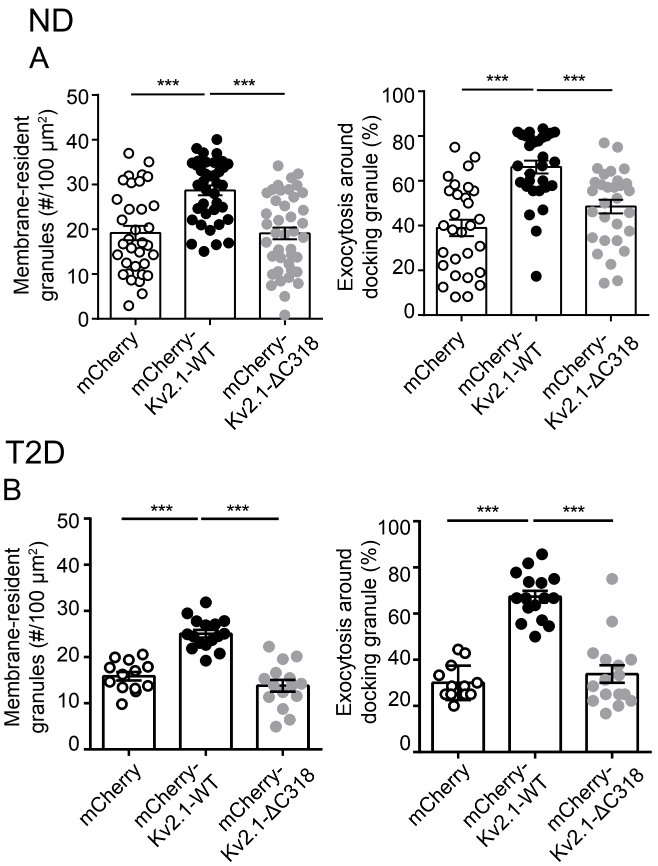

ND
A
***
***
Membrane-resident granules (#/100 µm²)
0
10
20
30
40
50
mCherry
mCherry-Kv2.1-WT
mCherry-Kv2.1-ΔC318
Exocytosis around docking granule (%)
0
20
40
60
80
100
T2D
B

**Figure.S3**

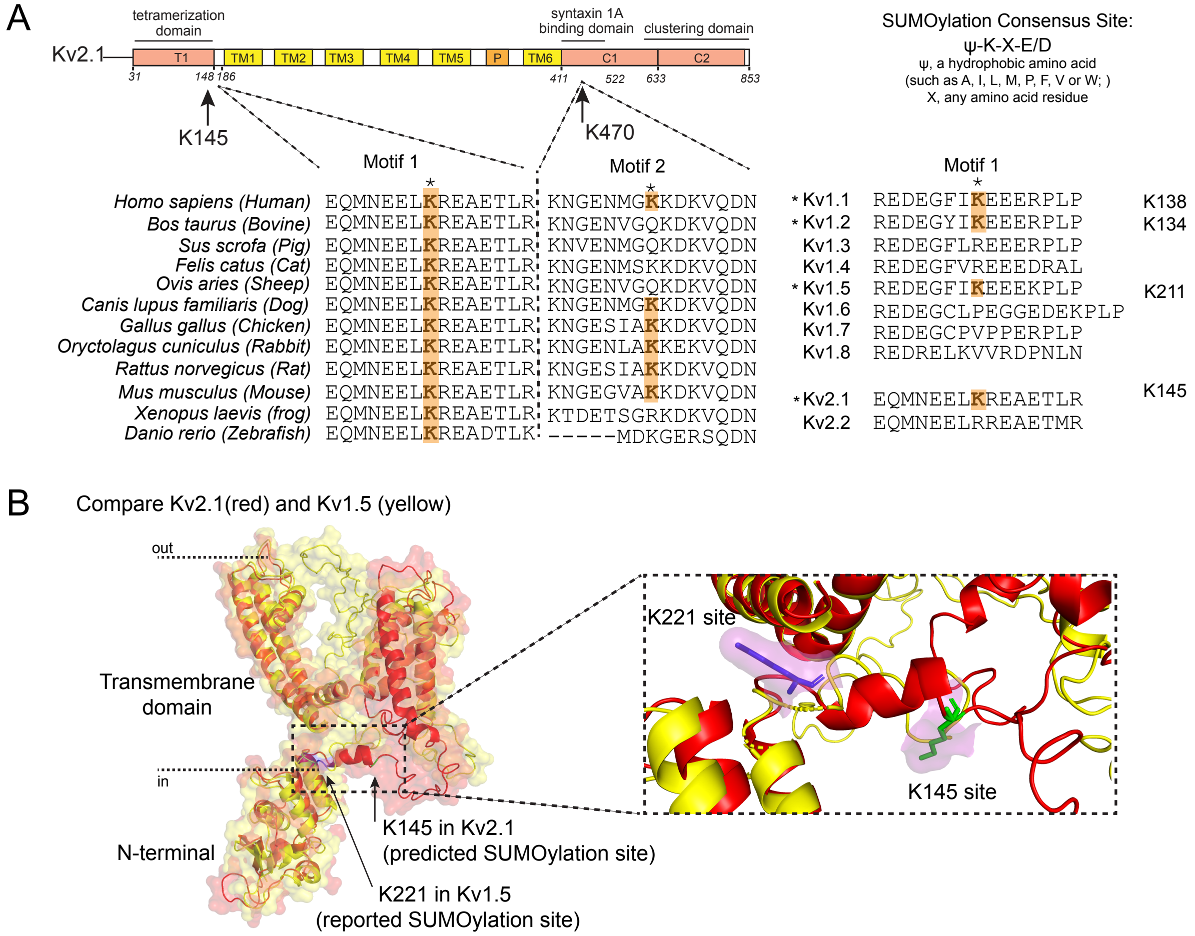
A
tetramerization domain
syntaxin 1A binding domain
clustering domain
Kv2.1
T1
TM1
TM2
TM3
TM4
TM5
P
TM6
C1
C2
31
148
186
411
522
633
853
K145
K470
Motif 1
Motif 2
Homo sapiens (Human) EQMNEELKREAETLR KNGENMGKKDKVQDN
Bos taurus (Bovine) EQMNEELKREAETLR KNGENVGQKDKVQDN
Sus scrofa (Pig) EQMNEELKREAETLR KNVENMGQKDKVQDN
Felis catus (Cat) EQMNEELKREAETLR KNGENMSKKDKVQDN
Ovis aries (Sheep) EQMNEELKREAETLR KNGENVGQKDKVQDN
Canis lupus familiaris (Dog) EQMNEELKREAETLR KNGENMGKKDKVQDN
Gallus gallus (Chicken) EQMNEELKREAETLR KNGESIAKKDKVQDN
Oryctolagus cuniculus (Rabbit) EQMNEELKREAETLR KNGENLAKKEKVQDN
Rattus norvegicus (Rat) EQMNEELKREAETLR KNGESIAKKDKVQDN
Mus musculus (Mouse) EQMNEELKREAETLR KNGEGVAKKDKVQDN
Xenopus laevis (frog) EQMNEELKREAETLR KTDETSGRKDKVQDN
Danio rerio (Zebrafish) EQMNEELKREADTLK -----MDKGERSQDN
SUMOylation Consensus Site:
ψ-K-X-E/D
ψ, a hydrophobic amino acid
(such as A, I, L, M, P, F, V or W; )
X, any amino acid residue
Motif 1
*Kv1.1 REDEGFIKEEERPLP K138
*Kv1.2 REDEGYIKEEERPLP K134
Kv1.3 REDEGFLREEERPLP
Kv1.4 REDEGFVREEEDRAL
*Kv1.5 REDEGFIKEEEKPLP K211
Kv1.6 REDEGCLPEGGEDEKPLP
Kv1.7 REDEGCPVPPERPLP
Kv1.8 REDRELKVVRDPNLN
*Kv2.1 EQMNEELKREAETLR K145
Kv2.2 EQMNEELRREAETMR
B
Compare Kv2.1(red) and Kv1.5 (yellow)
out
in
Transmembrane domain
N-terminal
K145 in Kv2.1
(predicted SUMOylation site)
K221 in Kv1.5
(reported SUMOylation site)
K221 site
K145 site

**Figure. S4**

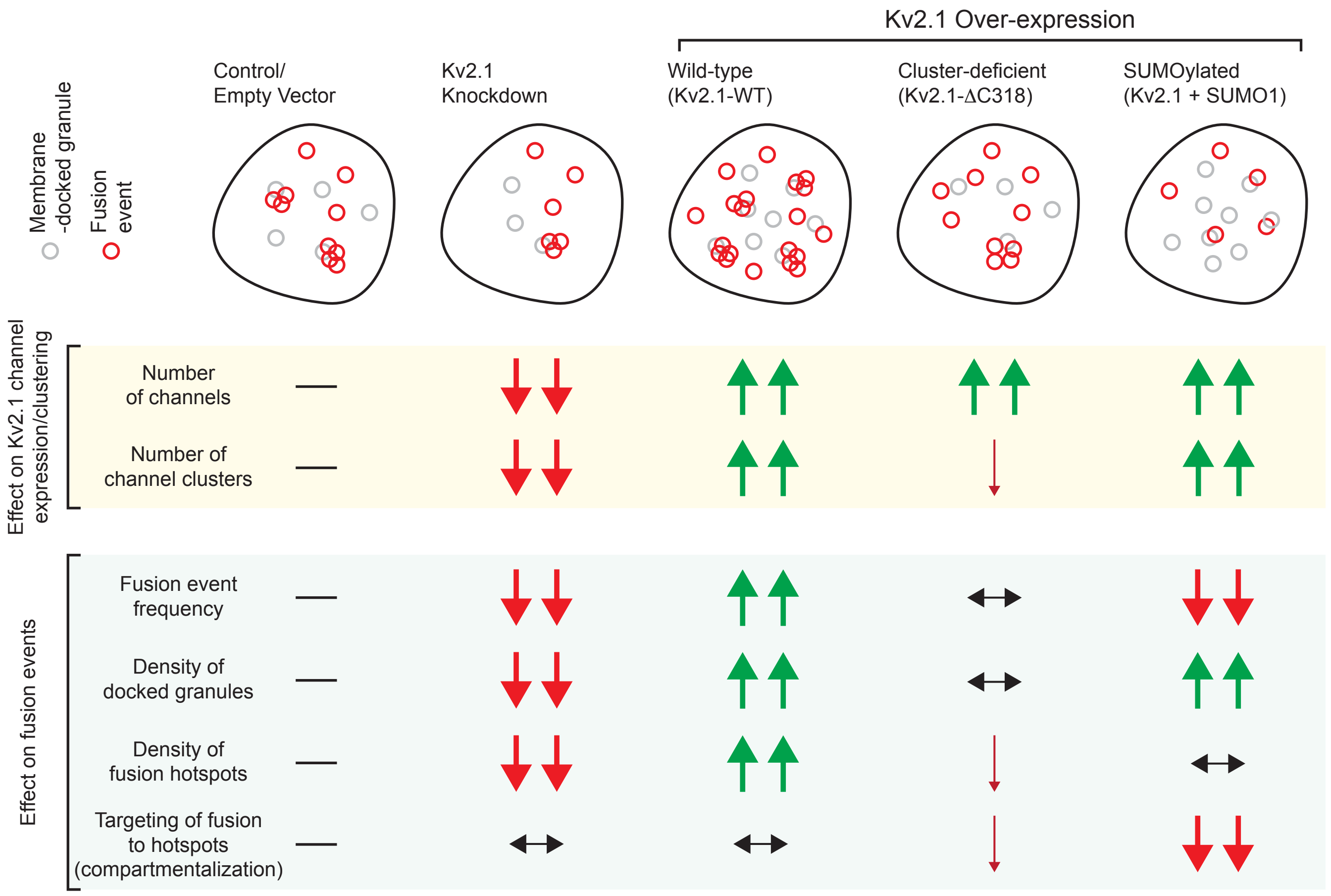

Kv2.1 Over-expression
Control/ Empty Vector
Kv2.1 Knockdown
Wild-type (Kv2.1-WT)
Cluster-deficient (Kv2.1-ΔC318)
SUMOylated (Kv2.1 + SUMO1)
Membrane -docked granule
Fusion event
Effect on Kv2.1 channel expression/clustering
Number of channels
Number of channel clusters
Effect on fusion events
Fusion event frequency
Density of docked granules
Density of fusion hotspots
Targeting of fusion to hotspots (compartmentalization)
